# Primary and secondary microplastics do not affect hatching of Japanese flounder eggs

**DOI:** 10.1101/2025.04.15.648670

**Authors:** Siti Syazwani Azmi, Taekyoung Seong, Hee-Jin Kim, Hisayuki Nakatani, Yusaku Kyozuka, Hiroshi Asakura, Kenichi Shimizu, Mitsuharu Yagi

## Abstract

Microplastics (MPs) are pervasive pollutants that may threaten aquatic organisms, especially during early life stages. This study investigated effects of MPs on the hatching rate of Japanese flounder eggs. Fertilized eggs were exposed to polystyrene (PS) microbeads (primary microplastics) (3 and 10 µm, at 20 and 200 particles/m*l*), and secondary MPs derived from coastal debris (rope, plastic bottles, fish net, string, and rubber pads) collected in Nagasaki, Japan. Hatching rates of flounder eggs were unaffected by either primary or secondary microplastics, suggesting limited impact of microplastics on this brief developmental stage.

## Introduction

Mismanagement of plastic waste has led to global plastic pollution estimated at around 8 million tons of plastic waste entering the ocean annually (Mallik et al. 2021). The term, “microplastics” (MPs), refers to plastic particles less than 5 mm in size (Du et al. 2021), which are categorized as primary or secondary microplastics, depending on their origin. Primary MPs are those originally produced as small particles, whereas secondary MPs result from degradation of larger plastic materials through chemical, physical, and biological processes (Cole et al. 2011). MPs are now found in almost every habitat, but predominantly occur in oceans, raising concerns about their potential impacts on marine life (Van Cauwenberghe et al. 2015; Auta et al. 2017; Cong et al. 2019; Borrelle et al. 2020). The most frequent interactions between marine organisms and MPs occur through ingestion, which has been observed in diverse groups of organisms, suggesting that MPs pose a threat to their survival (Cole et al. 2013; Desforges et al. 2015; Pedà et al. 2016; Jin et al. 2018).

Numerous studies have documented harmful impacts of MPs on development of embryos and early life stages of fish, including impaired feeding and DNA degradation (Japanese medaka) (de Sá et al. 2018; Pannetier et al. 2020), lower hatching and survival rates of zebrafish (Chen et al. 2017; Pitt et al. 2018; Santos et al. 2020), or even delayed hatching and growth of marine medaka (Li et al. 2020) under either high (Nobre et al. 2015; Gandara e Silva et al. 2016; Martínez-Gómez et al. 2017) or low concentrations of MPs (Li et al. 2020; Santos et al. 2020). Moreover, depending on their size, MPs can also cause physical obstruction of the digestive tract, leading to intestinal perforation, ulcers, and potentially gastric rupture, resulting in death (Law, 2017; Kim et al. 2021; Kim et al. 2022).

Embryonic development is a critical stage in the life cycles of fish. The chorion, a specialized structure enveloping the embryo until hatching, serves a protective function by impeding the ingress of various pollutants (Kristofco et al. 2018). Previous studies have revealed attachment of MPs to epithelia of zebrafish (*Danio rerio*) embryonic chorions (Batel et al. 2018), and attachment of plastic particles to chorions can create a localized hypoxic microenvironment, which may delay hatching (Zhang et al. 2021). Although fish are the most extensively studied taxonomic group in regard to MP effects (de Sá et al. 2018), there is little information concerning MP effects on embryos or the yolk-sac stage, which are highly sensitive to environmental and anthropogenic stressors (Pannetier et al. 2020; Uy and Johnson 2022).

Japanese flounder (*Paralichthys olivaceus*) is an economically important cultured fish in East Asia, particularly in Japan, Korea, and China, but very little literature has documented MP impacts on flounders. Wang et al. (2022) observed growth retardation of Japanese flounder under low (20 µg/*l*) or high (200 µg/*l*) concentrations of MPs, and Lee (2022) observed decreased growth and survival in starry flounder (*Platichthys stellatus*) under exposure to high concentrations of MPs. Therefore, the present study investigated whether exposure to primary and secondary MPs significantly affects hatching of Japanese flounder embryos at low and high MP concentrations.

## Materials and Methods

Fertilized eggs of Japanese flounder were purchased from Pacific Trading Co. LTD. After transport to the laboratory, eggs were acclimated for 4 h in a 30-L polycarbonate tank containing artificial seawater (33 ppt at 22 °C) before being used for experiments.

To investigate impacts of MP size and concentration on Japanese flounder, embryos (Experiment 1), commercial polystyrene (PS) microbeads (micromer®) of 3 and 10 µm diameter suspensions were purchased from micromod Partikeltechnologie GmbH, Schillingallee 68, D-18057 (Rostock, Germany). A few drops of PS stock suspensions were washed with distilled water in 50-mL centrifuge tubes, and these solutions were centrifuged at 9000 rpm for 10 min at 4 °C. Supernatants were removed and replaced with distilled water, and this procedure was repeated several times until a clear solution was obtained. These procedures were repeated to minimize effects of chemical substances possibly leaching from MPs during use. The final solution was diluted with sterilized seawater of the same salinity. MPs were counted under a microscope and the concentration was calculated before use.

Two MP stock solutions were prepared for 3-µm and 10 µm MPs. Each was diluted to final concentrations of 20 particles/m*l* and 200 particles/m*l* to yield four experimental groups plus controls (no MPs). All samples were prepared in triplicate.

To examine impacts of MP types on the hatching rate of Japanese flounder eggs (Experiment 2), marine coastal debris was collected at Shishigawa Fishing Port, Nagasaki, Japan (32° 51’ 49.8’’ N, 129° 48’ 22.6’’ E). Types of coastal debris were categorized as plastic bottles (PB), rope 1 (Ro1), rope 2 (Ro2), fishing line (FL), fishing net (FN), and tire fragments (TF). Coastal debris was washed with distilled water before freezing at - 80 °C. Then, then a collection of all coastal debris was ground into small particles using a household blender (Panasonic MK-K82) to yield micro-sized particles (MPs), which were then suspended in distilled water. These solutions were filtered through two cellulose ester membrane filters, first with a pore size of 3.00 µm and then of 1.00 µm (Advantec MFS Inc., Japan) to obtain purified solutions of 2.5-µm MPs, similar in size to phytoplankton (2 – 4 µm). For experimental trials, MP concentrations were set at 20 particles/m*l* for all coastal debris types.

MP suspensions (both polystyrene microbeads and coastal debris) were prepared in 33 ppt artificial seawater and distributed in 6-well plates (5 mL per well). Ten fertilized eggs, randomly selected from fish egg stocks, were added to the 6-well plates for all treatment groups, in triplicate. Cultures were agitated at 30 rpm using a shaker (NS-8 Neo Shaker, As ONE, Osaka, Japan) to prevent MP settling. Eggs were maintained at 22 °C under a 12L:12D photoperiod during the experiment. Daily observation was done thrice daily (9:00, 13:00, 17:00), and dead eggs and larvae were removed immediately. To maintain concentrations of MPs and water quality of the cultural medium in each well, surviving embryos and hatched larvae were transferred daily into new wells containing fresh MP suspension. Based on numbers of dead eggs, dead larvae, and hatched larvae, we calculated the hatching rate from the total number of fertilized eggs. Statistical analysis to determine MP impacts on the hatching rate of Japanese flounder eggs was performed using one-way analysis of variance (ANOVA). Before one-way ANOVA, Levene’s test was employed to verify homogeneity of variances. Subsequently, hatching rates of experimental groups were compared using Tukey and Duncan’s tests (*P* < 0.05). All statistical analyses were performed with Sigma Stat 3.0, SPSS, Chicago, U.S.A.

## Results

Results of Experiment 1 showed that the hatching rate of Japanese flounder was >90.0% in all experimental groups, including controls. All exposure groups except BH group showed hatching rates reduced by at least 5%, compared with controls (Fig.1), but no significant differences were observed among MP treatments (Tukey’s test, *P* > 0.05).

**Fig. 1.**
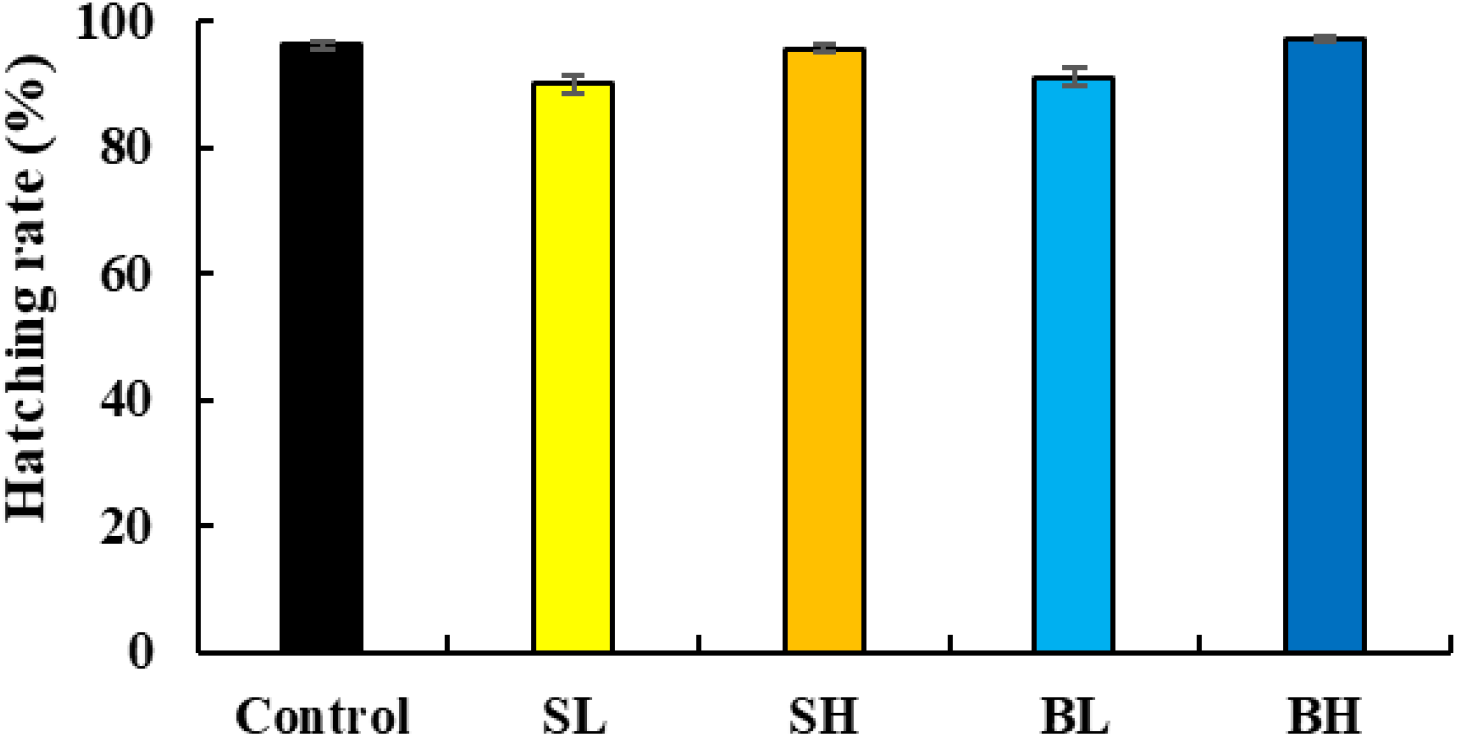
Hatching rate (%) of Japanese flounder under different microplastic concentrations 0 particles/m*l* (control), 3 µm/20 particles/m*l* (SL), 3 µm/200 particles/m*l* (SH), 10 µm/20 particles/m*l* (BL) and 10 µm/200 particles/m*l* (BH). Columns and error bars indicate means and standard deviations (n = 3).

Results of Experiment 2 showed that the hatching rate of Japanese flounder varied from 86.1% to 90.6% in all experimental groups and controls. All exposure groups showed hatching rates reduced by at least 5%, compared with controls (Fig. 2), but no significant differences were observed among treatments (Tukey’s test, *P* > 0.05).

**Fig. 2.**
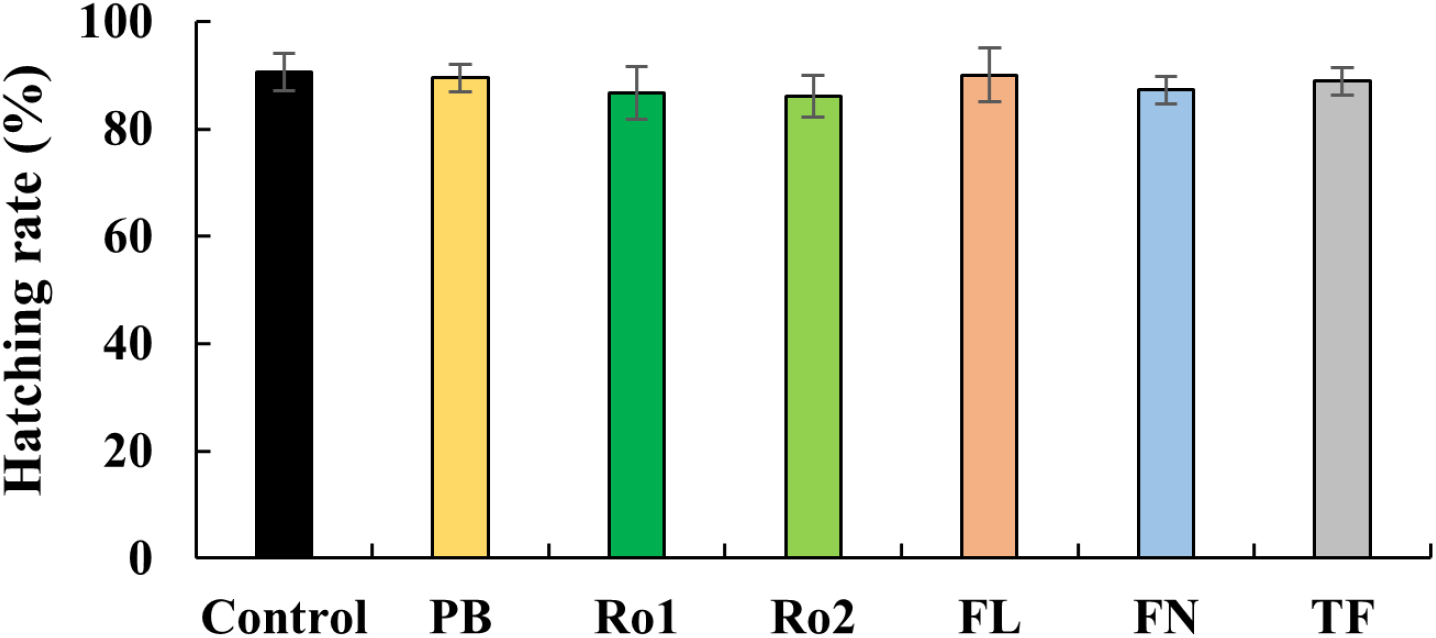
Hatching rate (%) of Japanese flounder expose to different types of secondary microplastics. Control, Plastic bottle (PB), Rope 1 (Ro1), Rope 2 (Ro2), Fishing line (FL), Fishing net (FN), and tire fragments (TF) under concentration of 20 particles/ml. Columns and error bars indicate means and standard deviations (n = 3).

## Discussion

The hatching rate of Japanese flounder eggs in both experiments exceeded 85% with both microbeads and coastal debris. It appeared not to be significantly affected by MPs of any size, type, or concentration. These results obtained are not entirely surprising, as embryos are considered well-protected from the surrounding environment by their chorions. However, there is a conceivable risk of small MPs accumulating on or adhering to eggs, potentially obstructing chorionic pores, as observed in other fish species, which has been linked to decreased hatching rates (Li et al. 2020; Malafaia et al. 2020). This phenomenon also may reflect to the 5% of decreasing hatching rates in the present study. Other fish studies have also reported that the presence of MPs did not correlate with hatching rate, as in three-spined sticklebacks (*Gasterosteus aculeatus*), zebrafish (*Danio rerio*) (LeMoine et al. 2018; Pitt et al. 2018; Jakubowska et al. 2020; Bunge (née Rebelein) et al. 2021), and marine medaka (*Oryzias melastigma*) (Beiras et al. 2018; Wang et al. 2021) exposed to MPs or even nanoplastics of various polymer types.

We speculate that the lack of significant effects of MPs in this experiment could be related to the size of MPs used. Most studies that reported negative effects on hatching rate employed plastic nanoparticles (<100 nm), which eventually penetrate the chorion, resulting in delayed hatching (Pitt et al. 2018; Duan et al. 2020). Micron-sized MPs are probably too large to cross the chorionic barrier, so the impact on embryos was negligible (Zhang et al. 2021). Nonetheless, the lack of significant effects observed in this study should not be interpreted as evidence that MPs are universally safe for aquatic organisms regardless of their biology. These conflicting results highlight the complexity of MP pollution and emphasize the necessity for more extensive investigations. Future research should examine longer-term effects of MPs on aquatic organisms, as some impacts may manifest themselves chronically rather than acutely (Huerta Lwanga et al. 2016; Naidoo and Glassom 2019). Additionally, it is essential to explore how MPs affect different life stages of organisms, considering that vulnerabilities and responses to environmental stressors can vary significantly during development (LeMoine et al. 2018; Pitt et al. 2018; Li et al. 2020).

## Conclusion

In conclusion, our findings reveal that neither primary nor secondary MPs negatively impact the hatching rate of Japanese flounder eggs, as more than 85% successfully hatched into larvae. However, more comprehensive research is needed to grasp the full implications of MPs on aquatic organisms. Future MP research should assess growth, development, and survival of marine organisms in various life stages, including interactions with other stressors, in order to assess broader effects of MP pollution on aquatic organisms.

## Acknowledgements

This research was supported by the Environment Research and Technology Development Fund (1MF-2204) of the Environmental Restoration and Conservation Agency provided by the Ministry of Environment of Japan.

## References

Auta, H. S., Emenike, C. U. and Fauziah, S. H. (2017). Distribution and importance of microplastics in the marine environment: A review of the sources, fate, effects, and potential solutions. Environ. Int., 102, 165–176. 10.1016/j.envint.2017.02.013

Batel, A., Borchert, F., Reinwald, H., Erdinger, L. and Braunbeck, T. (2018). Microplastic accumulation patterns and transfer of benzo[a]pyrene to adult zebrafish (Danio rerio) gills and zebrafish embryos. Environ. Pollut., 235, 918–930. 10.1016/j.envpol.2018.01.028

Beiras, R., Bellas, J., Cachot, J., Cormier, B., Cousin, X., Engwall, M., Gambardella, C., Garaventa, F., Keiter, S., Le Bihanic, F., López-Ibáñez, S., Piazza, V., Rial, D., Tato, T. and Vidal-Liñán, L. (2018). Ingestion and contact with polyethylene microplastics does not cause acute toxicity on marine zooplankton. J. of Hazard. Mater., 360, 452–460. 10.1016/j.jhazmat.2018.07.101

Borrelle, S. B., Ringma, J., Law, K. L., Monnahan, C. C., Lebreton, L., McGivern, A., Murphy, E., Jambeck, J., Leonard, G. H., Hilleary, M. A., Eriksen, M., Possingham, H. P., De Frond, H., Gerber, L. R., Polidoro, B., Tahir, A., Bernard, M., Mallos, N., Barnes, M. and Rochman, C. M. (2020). Predicted growth in plastic waste exceeds efforts to mitigate plastic pollution. Sci, 369(6510), 1515– 1518. 10.1126/science.aba3656

Bunge (née Rebelein), A., Kammann, U. and Scharsack, J. P. (2021). Exposure to microplastic fibers does not change fish early life stage development of three-spined sticklebacks (Gasterosteus aculeatus). Microplastics and Nanoplastics, 1(1), 15. 10.1186/s43591-021-00015-x

Chen, Q., Gundlach, M., Yang, S., Jiang, J., Velki, M., Yin, D. and Hollert, H. (2017). Quantitative investigation of the mechanisms of microplastics and nanoplastics toward zebrafish larvae locomotor activity. Sci. Total Environ., 584–585, 1022– 1031. 10.1016/j.scitotenv.2017.01.156

Cole, M., Lindeque, P., Halsband, C. and Galloway, T. S. (2011). Microplastics as contaminants in the marine environment: A review. Mar. Pollut. Bull., 62(12), 2588–2597. 10.1016/j.marpolbul.2011.09.025

Cole, M., Lindeque, P., Fileman, E., Halsband, C., Goodhead, R., Moger, J. and Galloway, T. S. (2013). Microplastic Ingestion by Zooplankton. Environ. Sci. & Technol., 47(12), 6646–6655. 10.1021/es400663f

Cong, Y., Jin, F., Tian, M., Wang, J., Shi, H., Wang, Y. and Mu, J. (2019). Ingestion, egestion and post-exposure effects of polystyrene microspheres on marine medaka (Oryzias melastigma). Chemosphere, 228, 93–100. 10.1016/j.chemosphere.2019.04.098

de Sá, L. C., Oliveira, M., Ribeiro, F., Rocha, T. L. and Futter, M. N. (2018). Studies of the effects of microplastics on aquatic organisms: What do we know and where should we focus our efforts in the future? Sci. Total Environ., 645, 1029–1039. 10.1016/j.scitotenv.2018.07.207

Desforges, J. P. W., Galbraith, M. and Ross, P. S. (2015). Ingestion of Microplastics by Zooplankton in the Northeast Pacific Ocean. Arch. Environ. Contam. and Toxicol., 69(3), 320–330. 10.1007/s00244-015-0172-5

Du, S., Zhu, R., Cai, Y., Xu, N., Yap, P.-S., Zhang, Y., He, Y. and Zhang, Y. (2021). Environmental fate and impacts of microplastics in aquatic ecosystems: A review. RSC Advances, 11(26), 15762–15784. 10.1039/D1RA00880C

Duan, Z., Duan, X., Zhao, S., Wang, X., Wang, J., Liu, Y. and Wang, L. (2020). Barrier function of zebrafish embryonic chorions against microplastics and nanoplastics and its impact on embryo development. J. of Hazard. Mater., 395, 122621.

Gandara e Silva, P. P., Nobre, C. R., Resaffe, P., Pereira, C. D. S. and Gusmão, F. (2016). Leachate from microplastics impairs larval development in brown mussels. Water Res., 106, 364–370. 10.1016/j.watres.2016.10.016

Huerta Lwanga, E., Gertsen, H., Gooren, H., Peters, P., Salánki, T., van der Ploeg, M., Besseling, E., Koelmans, A. A. and Geissen, V. (2016). Microplastics in the Terrestrial Ecosystem: Implications for Lumbricus terrestris (Oligochaeta, Lumbricidae). Environ. Sci. & Technol., 50(5), 2685–2691. 10.1021/acs.est.5b05478

Jakubowska, M., Białowąs, M., Stankevičiūtė, M., Chomiczewska, A., Pažusienė, J., Jonko-Sobuś, K., Hallmann, A. and Urban-Malinga, B. (2020). Effects of chronic exposure to microplastics of different polymer types on early life stages of sea trout Salmo trutta. Sci. Total Environ., 740, 139922. 10.1016/j.scitotenv.2020.139922

Jin, Y., Xia, J., Pan, Z., Yang, J., Wang, W. and Fu, Z. (2018). Polystyrene microplastics induce microbiota dysbiosis and inflammation in the gut of adult zebrafish. Environ. Pollut., 235, 322–329. 10.1016/j.envpol.2017.12.088

Kim, J. H., Yu, Y. B. and Choi, J. H. (2021). Toxic effects on bioaccumulation, hematological parameters, oxidative stress, immune responses and neurotoxicity in fish exposed to microplastics: A review. J. of Hazard. Mater., 413, 125423. 10.1016/j.jhazmat.2021.125423

Kim, S. A., Kim, L., Kim, T. H. and An, Y. J. (2022). Assessing the size-dependent effects of microplastics on zebrafish larvae through fish lateral line system and gut damage. Mar. Pollut. Bull., 185, 114279. 10.1016/j.marpolbul.2022.114279

Kristofco, L. A., Haddad, S. P., Chambliss, C. K. and Brooks, B. W. (2018). Differential uptake of and sensitivity to diphenhydramine in embryonic and larval zebrafish. Environ. Toxicol. and Chem., 37(4), 1175–1181. 10.1002/etc.4068

Law, K. L. (2017). Plastics in the Marine Environment. Annu. Rev. of Mar. Sci., 9(Volume 9, 2017), 205–229. 10.1146/annurev-marine-010816-060409

Lee J. H. (2022). Responses of hematological parameters, antioxidant and heat shock protein 90 in starry flounder, Platichthys stellatus exposed to microplastics. https://repository.pknu.ac.kr:8443/handle/2021.oak/32714

LeMoine, C. M. R., Kelleher, B. M., Lagarde, R., Northam, C., Elebute, O. O. and Cassone, B. J. (2018). Transcriptional effects of polyethylene microplastics ingestion in developing zebrafish (Danio rerio). Environ. Pollut., 243, 591–600. 10.1016/j.envpol.2018.08.084

Li, Y., Wang, J., Yang, G., Lu, L., Zheng, Y., Zhang, Q., Zhang, X., Tian, H., Wang, W. and Ru, S. (2020). Low level of polystyrene microplastics decreases early developmental toxicity of phenanthrene on marine medaka (Oryzias melastigma). J. of Hazard. Mater., 385, 121586. 10.1016/j.jhazmat.2019.121586

Malafaia, G., de Souza, A. M., Pereira, A. C., Gonçalves, S., da Costa Araújo, A. P., Ribeiro, R. X. and Rocha, T. L. (2020). Developmental toxicity in zebrafish exposed to polyethylene microplastics under static and semi-static aquatic systems. Sci. Total Environ., 700, 134867. 10.1016/j.scitotenv.2019.134867

Mallik, A., Xavier, K. A. M., Naidu, B. C. and Nayak, B. B. (2021). Ecotoxicological and physiological risks of microplastics on fish and their possible mitigation measures. Sci. Total Environ., 779, 146433. 10.1016/j.scitotenv.2021.146433

Martínez-Gómez, C., León, V. M., Calles, S., Gomáriz-Olcina, M. and Vethaak, A. D. (2017). The adverse effects of virgin microplastics on the fertilization and larval development of sea urchins. Mar. Environ. Res., 130, 69–76. 10.1016/j.marenvres.2017.06.016

Naidoo, T. and Glassom, D. (2019). Decreased growth and survival in small juvenile fish, after chronic exposure to environmentally relevant concentrations of microplastic. Mar. Pollut. Bull., 145, 254–259. 10.1016/j.marpolbul.2019.02.037

Nobre, C. R., Santana, M. F. M., Maluf, A., Cortez, F. S., Cesar, A., Pereira, C. D. S. and Turra, A. (2015). Assessment of microplastic toxicity to embryonic development of the sea urchin Lytechinus variegatus (Echinodermata: Echinoidea). Mar. Pollut. Bull., 92(1), 99–104. 10.1016/j.marpolbul.2014.12.050

Pannetier, P., Morin, B., Le Bihanic, F., Dubreil, L., Clérandeau, C., Chouvellon, F., Van Arkel, K., Danion, M. and Cachot, J. (2020). Environmental samples of microplastics induce significant toxic effects in fish larvae. Environ. Internat., 134, 105047. 10.1016/j.envint.2019.105047

Pedà, C., Caccamo, L., Fossi, M. C., Gai, F., Andaloro, F., Genovese, L., Perdichizzi, A., Romeo, T. and Maricchiolo, G. (2016). Intestinal alterations in European sea bass Dicentrarchus labrax (Linnaeus, 1758) exposed to microplastics: Preliminary results. Environ. Pollut., 212, 251–256. 10.1016/j.envpol.2016.01.083

Pitt, J. A., Kozal, J. S., Jayasundara, N., Massarsky, A., Trevisan, R., Geitner, N., Wiesner, M., Levin, E. D. and Di Giulio, R. T. (2018). Uptake, tissue distribution, and toxicity of polystyrene nanoparticles in developing zebrafish (Danio rerio). Aquat. Toxicol., 194, 185–194. 10.1016/j.aquatox.2017.11.017

Pitt, J. A., Trevisan, R., Massarsky, A., Kozal, J. S., Levin, E. D. and Di Giulio, R. T. (2018). Maternal transfer of nanoplastics to offspring in zebrafish (Danio rerio): A case study with nanopolystyrene. Sci. Total Environ., 643, 324–334. 10.1016/j.scitotenv.2018.06.186

Santos, D., Félix, L., Luzio, A., Parra, S., Cabecinha, E., Bellas, J. and Monteiro, S. M. (2020). Toxicological effects induced on early life stages of zebrafish (Danio rerio) after an acute exposure to microplastics alone or co-exposed with copper. Chemosphere, 261, 127748. 10.1016/j.chemosphere.2020.127748

Uy, C. A. and Johnson, D. W. (2022). Effects of microplastics on the feeding rates of larvae of a coastal fish: Direct consumption, trophic transfer, and effects on growth and survival. Mar. Biol., 169(2), 27. 10.1007/s00227-02104010-x

Van Cauwenberghe, L., Devriese, L., Galgani, F., Robbens, J. and Janssen, C. R. (2015). Microplastics in sediments: A review of techniques, occurrence and effects. Mar. Environ. Res., 111, 5–17. 10.1016/j.marenvres.2015.06.007

Wang, J., Zheng, M., Lu, L., Li, X., Zhang, Z. and Ru, S. (2021). Adaptation of lifehistory traits and trade-offs in marine medaka (Oryzias melastigma) after whole life-cycle exposure to polystyrene microplastics. J. of Hazard. Mater., 414, 125537. 10.1016/j.jhazmat.2021.125537

Wang, J., Li, X., Li, P., Li, L., Zhao, L., Ru, S. and Zhang, D. (2022). Porous microplastics enhance polychlorinated biphenyls-induced thyroid disruption in juvenile Japanese flounder (Paralichthys olivaceus). Mar. Pollut. Bull., 174, 113289. 10.1016/j.marpolbul.2021.113289

Zhang, C., Wang, J., Zhou, A., Ye, Q., Feng, Y., Wang, Z., Wang, S., Xu, G. and Zou, J. (2021). Species-specific effect of microplastics on fish embryos and observation of toxicity kinetics in larvae. J. of Hazard. Mater., 403, 123948. 10.1016/j.jhazmat.2020.1239

